# Intragenomic Redistribution of Host Transcription Factor Binding With *Toxoplasma Gondii* Infection

**DOI:** 10.1101/132076

**Authors:** Netha Ulahannan, Masako Suzuki, Claudia A. Simões-Pires, Zofia Wicik, N. Ari Wijetunga, Matthew McKnight Croken, Sanchari Bhattacharyya, Andrew D. Johnston, Yu Kong, Shahina B. Maqbool, Amit Verma, John M. Greally

**Affiliations:** Department of Microbiology and Immunology, Albert Einstein College of Medicine, Bronx, NY 10461; Department of Genetics, Albert Einstein College of Medicine, Bronx, NY 10461; Department of Developmental and Molecular Biology, Albert Einstein College of Medicine, Bronx, NY 10461; Department of Medicine, Albert Einstein College of Medicine, Bronx, NY 10461

## Abstract

The intracellular pathogen *Toxoplasma gondii* modifies a number of host cell processes. The mechanisms by which *T. gondii* alters host gene expression are incompletely understood. This study focuses on how the regulators of gene expression in human host cells respond to *T. gondii* 24 hours following infection to cause specific patterns of transcriptional dysregulation. The most striking finding was the altered landscape of transposase-accessible chromatin by infection. We found both gains and losses of loci of open chromatin enriched in proximity to transcriptionally altered genes. Both DNA sequence motif analysis at the loci changing chromatin accessibility and network analysis of the genes with transcription and regulatory changes implicate a central role for the AP-1 transcription factor. We validated the redistribution of AP-1 in the host genome using chromatin immunoprecipitation studies of the c-Fos component of AP-1. As infection with *T. gondii* is associated with the cell failing to progress through the cell cycle, all of the changes observed occur in the absence of cell division and within 24 hours, an insight into the dynamism of these transcriptional regulatory events. We conclude that *T. gondii* infection influences transcriptional regulation through transcription factor re-targeting to modify the *cis*-regulatory landscape of the host nucleus.

**AUTHOR SUMMARY:** The complex interactions between the intracellular pathogen *Toxoplasma gondii* and the host cell manifest as expression changes of host genes. *T. gondii’s* secreted effectors have been extensively studied and include factors that influence the properties of transcription factors, resulting in post-translational modifications and changes in intracellular localization. To gain insights into how *T. gondii* exerts specific influences on host transcriptional regulation, we used genome-wide approaches to study gene expression, cytosine modifications, and chromatin structure of the host cell 24 hours after infection. The greatest insights were gained from the mapping of loci of transposase-accessible chromatin, revealing a consistently altered pattern of a subset of loci becoming inaccessible, with the simultaneous acquisition of a new set of infection-associated loci of open chromatin. The sequences at these loci were enriched for certain transcription factor binding motifs, in particular that of AP-1, the transcription factor formed by c-Jun and c-Fos heterodimers. Network analysis revealed a central role for c-Jun and c-Fos in the infection-associated perturbations, prompting a chromatin immunoprecipitation approach that confirmed the redistribution of c-Fos in infected cells. We conclude that a *T. gondii* infection leads to an intragenomic redistribution of host transcription factor binding, with resulting effects on host gene expression.

## INTRODUCTION

Cellular responses to perturbations can involve transcriptional alterations, which can be transient, or can involve residual, long-term reprogramming events that are defined as epigenetic mechanisms of cellular memory. Intracellular infections represent a relatively tractable model of host cell perturbation, with characteristic transcriptional alterations that have prompted numerous studies of potential regulatory modifications that may be mediating these changes. Acute *in vitro* bacterial infection of human leukocytes by *Mycobacterium tuberculosis* [1,2], *Anaplasma phagocytophilum* [3], *Burkholderia pseudomallei* [4] or *Leishmania donovani* [5] have been associated with DNA methylation changes of the host genome. Chromatin modifications of the host have also been characterized, with *Listeria monocytogenes* infection of human cell lines *in vitro* causing deacetylation of lysine 18 of histone H3 (H3K18) [6]. This process is initiated by the *Listeria* protein InIB, which activates PI3K and Akt signaling to induce nuclear translocation of the histone deacetylase sirtuin 2 (SIRT2), which in turn leads to the chromatin effects observed [7]. A separate process was described for the *Listeria monocytogenes* protein LntA (listeria nuclear targeted protein A), which interacts strongly with the host heterochromatin protein BAHD1 (bromo adjacent homology domain-containing 1 protein), causing it to be re-targeted within the genome to silence interferon (IFN)–stimulated genes [8]. Apart from these bacterial pathogens, there are also examples of eukaryotic intracellular parasites with effects on host transcriptional regulation. Infection of mice with *Plasmodium berghei* causes a transcriptional response in liver within 24 hours [9], with another apicomplexan parasite *Toxoplasma gondii* also found to induce altered host gene expression patterns [10–15]. Recent insights into the possible mechanisms of this transcriptional dysregulation include the finding that *T. gondii* secretes a protein called TgIST (*T. gondii* inhibitor of STAT1 transcriptional activity) that not only binds to and influences the genomic binding pattern of STAT1, it also recruits histone deacetylases and the nucleosome remodeling deacetylase (NuRD) transcriptional repressor complex [16,17]. The model that emerges from all of these examples is one of factors from intracellular pathogens binding to or causing post-translational modifications of host transcriptional regulators, resulting in alterations of the host’s transcriptional patterns.

The downstream effects of these infection-induced perturbations are very heterogeneous molecular processes. With the development of genome-wide assays, we are not restricted to a focus on specific genomic loci, and can explore both chromatin structure and DNA methylation as candidates for mediating transcriptional dysregulatory influences. We therefore designed a set of experiments studying these molecular mechanisms to test how the acute effects of *T. gondii* infection influence transcriptional regulatory processes in human cells.

## RESULTS

### Gene expression is altered in T. gondii-infected cells

We provide an overview of the experiments performed in **Figure S1**. We studied RH strain parasites in the tachyzoite stage of the *T. gondii* life cycle, infecting human foreskin fibroblasts (HFFs). The foundation for the project was a set of transcriptional studies, so that we could compare our results with those of prior studies to define the common transcriptional dysregulation patterns during a *T. gondii* infection. RNA was harvested 24 hours after infection from three sets each of uninfected and infected cells. RNA-seq performed on these samples revealed a number of host genes significantly changing expression with infection (Figure 1a), as expected from prior studies using microarrays [10–14,18]. An over-representation analysis (ORA) [19] of the dysregulated genes revealed enrichment for a number of functional pathways. These pathways overlap considerably with those identified in the preceding microarray-based gene expression studies of *T. gondii*-infected HFFs [10–14,18] as we show in **Table S1**, including metabolic (hypoxia, glycolysis), cell cycle and inflammatory pathways, with TNF alpha and IL2/STAT5 signaling and the targets of the MYC transcriptional regulator [10–14,18] especially prominent and concordant between studies. The 24 hour timepoint following *T. gondii* infection is therefore generating a pattern of transcriptional changes that reproduces those found in prior, separate studies at the same timepoint.

**Figure 1.**
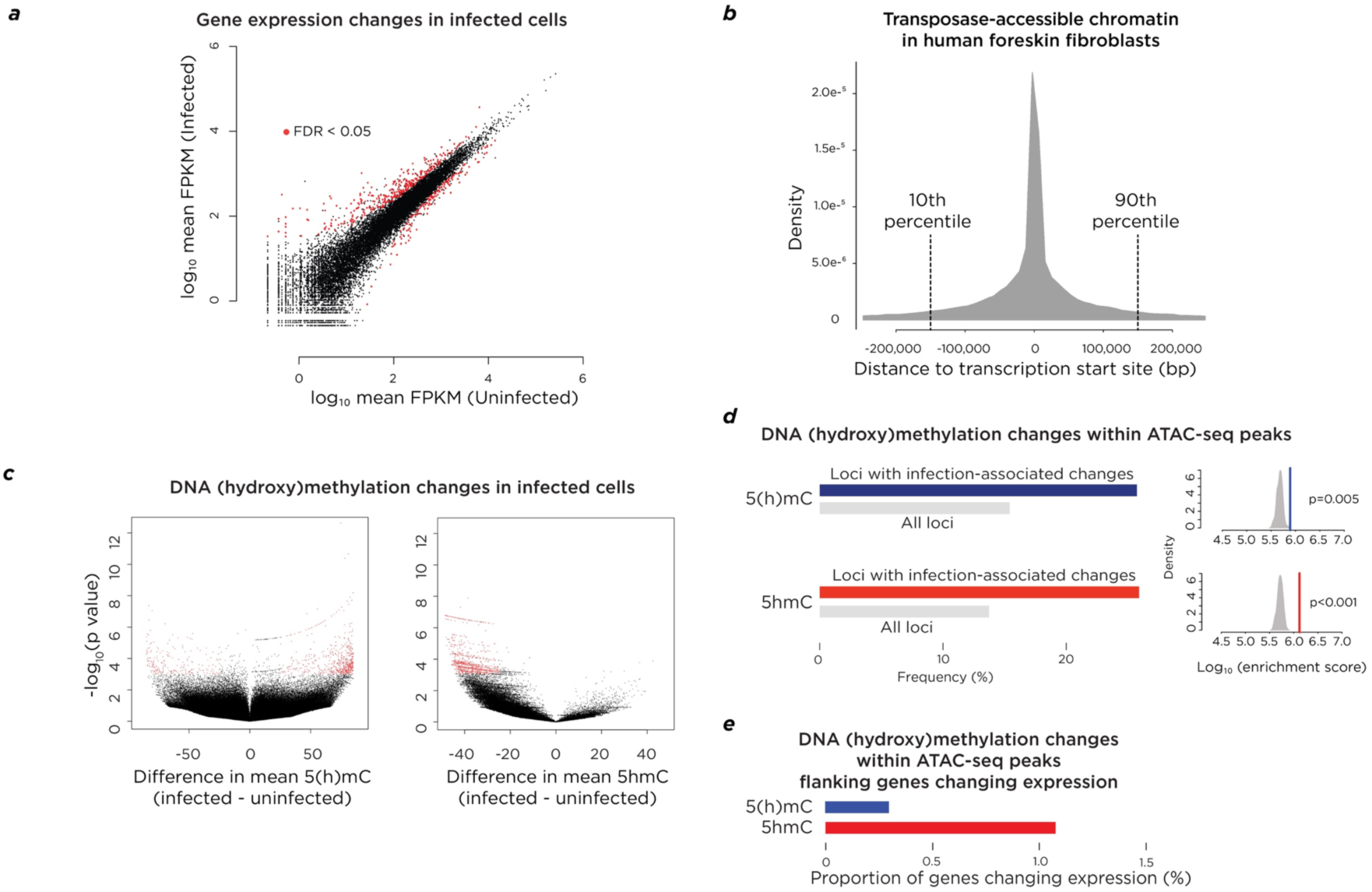
Changes in cytosine modifications are enriched near genes changing expression in cells infected with *T. gondii*. In **(a)** we show genes changing expression significantly in infected cells (red). In **(b)** we show that all but the most outlying upstream and downstream 10% of ATAC-seq peaks (indicating transposase-accessible chromatin) occur within a ∼150 kb region flanking annotated RefSeq gene transcription start sites (TSSs). In **(c)** we show volcano plots of the HELP-tagging and HELP-GT results, with the most significantly altered loci shown in red, and a prominent skewing in the HELP-GT plot consistent with a loss of 5hmC with infection. Panel **(d)** shows the enrichment of the proportion of loci tested that change 5(h)mC or 5hmC within the ATAC-seq peaks in the 150 kb flanking TSSs compared with genome-wide, with permutation plots showing this enrichment to be significant. The final panel **(e)** shows that the proportion of differentially-expressed genes with altered 5(h)mC or 5hmC within flanking ATAC-seq peaks is very low (∼1% for 5hmC), showing that the mechanism of transcriptional perturbation of most genes is not revealed by this combination of genome-wide approaches.

### *DNA methylation and hydroxymethylation alterations are enriched at* cis*-regulatory sites flanking a minority of genes changing expression*

To test how DNA methylation is altered during a *T. gondii* infection, we used the HELP-tagging assay [20], which measures cytosine methylation at up to 2 million HpaII sites throughout the human genome, testing more distal *cis*-regulatory elements than other commonly-used DNA methylation survey approaches [21]. The DNA harvested from the samples tested by expression profiling was used to generate the HELP-tagging libraries. The values obtained for each locus represents the combination of 5-methylcytosine (5mC) and a variable but generally much smaller proportion of 5-hydroxymethylcytosine (5hmC). We therefore refer to the information from HELP-tagging as a measure of 5(h)mC. We also performed the HELP-GT assay to quantify 5hmC at HpaII sites [22] on the same DNA samples.

To perform the analysis of changes associated with transcriptional dysregulation, we focused on changes at *cis*-regulatory elements. It is now well-established that the DNA methylation changes associated with cancer [23–26], non-cancerous conditions [27,28] and normal cell differentiation [29,30] occur more frequently at distal *cis*-regulatory elements other than promoters of genes. One such example is the “shores” flanking CpG islands, which dynamically change DNA methylation associated with local transcriptional alterations [31], loci that we have found to have the characteristics of transcriptional enhancers [32]. We have previously found enhancers to be the most informative loci for DNA methylation changes when studying renal fibrosis [27], suggesting that a similar approach in the HFFs infected by *T. gondii* would be of value. Rather than relying on annotations from public databases of comparable cell types, we used the assay for transposase-accessible chromatin using sequencing (ATAC-seq) [33] to define the *cis*-regulatory landscape in the HFFs we were infecting. The statistically-significant changes of DNA methylation within these loci were the focus for our analyses. To link DNA methylation changes to genes, we plotted the distribution of ATAC-seq peaks (representing open chromatin) centered on the transcription start site of each annotated RefSeq gene in the human genome. We found that all but the most distant upstream and downstream 10% of transposase-accessible loci are within ±150 kb flanking the transcription start sites (TSSs) of RefSeq genes (Figure 1b). When we tested the loci with the most significantly changed 5(h)mC and 5hmC (Figure 1c), we found that they were significantly enriched in these loci of transposase-accessible chromatin flanking TSSs (Figure 1d), but that only ∼1% or fewer of genes changing expression had 5(h)mC or 5hmC changes at flanking *cis*-regulatory sites (Figure 1e). While our findings are consistent with prior studies of intracellular infections that have been associated with DNA methylation changes of the host cell’s genome [1–5], they do not give us insight into how the majority of genes in the genome are transcriptionally perturbed.

### Increased aerobic glycolysis occurs with infection

One incidental finding from the HELP-GT data was that of a global shift towards decreased 5hmC in the infected cells (Figure 1c). Decreased 5hmC globally could be due to decreased activity of the TET enzymes that convert 5mC to 5hmC, which in turn may be linked to changes in cofactor availability that occur when cells alter their metabolic state. We therefore tested expression levels of genes associated with glycolysis (*LDHA, PDK1, GLUT1*), finding them all to be significantly up-regulated in expression (Figure 2a). This indicates a shift to aerobic glycolysis in these acutely infected cells, possibly related to *T. gondii’s* known effect on host HIF1alpha [34]. As TET enzymes are dependent on iron and alpha-ketoglutarate as co-factors [35], a change in the metabolic state of the cell could potentially induce changes in TET activity. There was no significant change in the levels of transcripts for *TET1-TET3* (Figure 2b), nor was there a change in the amount of TET activity when adjusted by cell number (Figure 2c), which would otherwise be misinterpreted if measured as a function of total protein concentration, as the infected cells contain *T. gondii* proteins not present in the uninfected cells. In Figure 2d we show the results of mass spectrometry for 5mC and 5hmC in uninfected and infected cells. We have previously shown that *T. gondii* has no 5(h)mC [36], so the DNA sample from the infected cells could be artefactually skewed towards decreased 5mC/5hmC. Even with this caveat, the mass spectrometry results show no significant changes of 5mC or 5hmC with infection. The illustration of the very small proportion of 5hmC relative to 5mC (Figure 2e) indicates why a global shift in 5hmC in the HELP-GT assay is not reflected by some degree of loss of 5(h)mC in the HELP-tagging assay (Figure 1c). We conclude that while the subset of loci tested by the HELP-GT assay show a trend towards the loss of 5hmC, our further validation studies do not support these reflect changes in the overall levels of 5hmC in the cell, nor an effect upon TET activity due to the shift in infected cells towards aerobic glycolysis.

**Figure 2.**
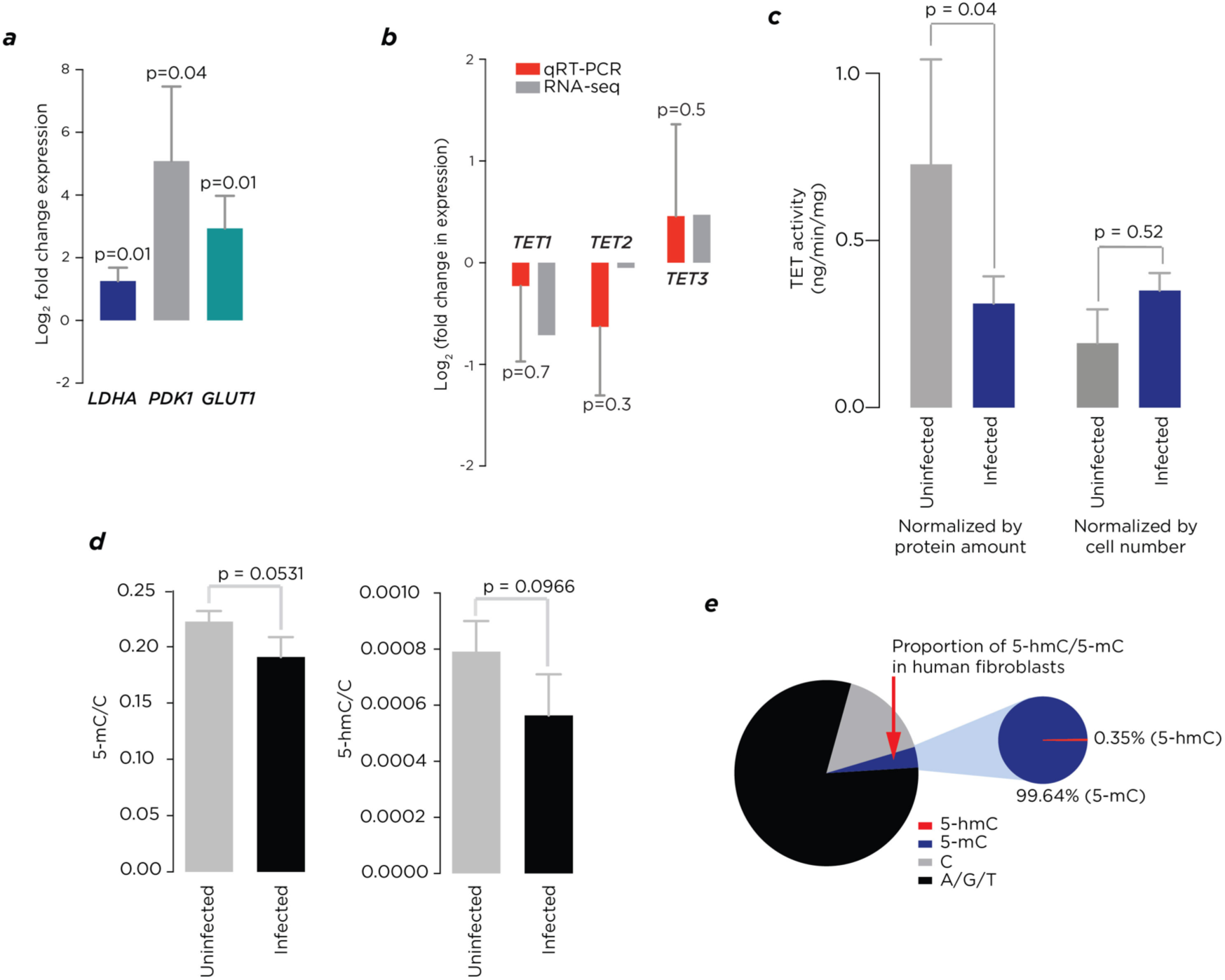
*T. gondii* infection is associated with a switch to aerobic glycolysis in infected host cells. The initial observation was from the volcano plot in Figure 1 that showed an unexpected skewing indicating the loss of 5hmC in infected cells. To test whether this reflected an altered metabolic state **(a)**, we tested the expression levels of *LDHA, PDK1* and *GLUT1*, their concordant significant increases in expression supporting the switch to increased aerobic glycolysis in the infected cells. This was not accompanied by any changes in expression levels of *TET* genes **(b)**, and while overall TET enzyme activity was significantly lower in protein extracted from nuclei from infected cells **(c)**, the infected cells have a contribution of protein from *T. gondii*, requiring that we normalize by host cell number, which eliminates the difference. Mass spectroscopy was performed to test the 5mC and 5hmC changes with infection throughout the genome, finding no significant changes **(d)**. This quantification allows us to depict the relative contributions of C, 5mC and 5hmC **(e)**, illustrating why even substantial changes in 5hmC are not reflected by changes in 5(h)mC.

### Loci with altered chromatin accessibility are enriched flanking genes changing expression

The approach of using a reference epigenome to allow a focus on *cis*-regulatory sequences makes the assumption that the locations of the regulatory elements remain unaltered with the cellular perturbation being studied. We therefore explored this question, adding ATAC-seq studies of the infected cells. We performed assays on different samples of uninfected and infected cells in triplicate. We mapped loci of open chromatin and categorized them into three groups of loci: those present in all six replicates (constitutive), those present only in all three replicates of uninfected cells (loci losing transposase-accessibility with infection) and those that only appeared in all three replicates of infected cells (loci gaining transposase accessibility with infection). The same ±150 kb region flanking RefSeq transcription start sites was found to contain all but the outlying 20% of ATAC-seq peaks gained or lost with infection (**Figure S2**). In total there were 76,420 constitutive loci, with 6,249 uninfected-only loci and 4,096 infected-only loci. Therefore, of the 82,669 loci of open chromatin mapped in uninfected cells, 7.6% are lost with infection, with the gain of a proportion of new loci of open chromatin equivalent to 5.0% of the pre-existing loci. The reference epigenome approach is therefore found to be a reasonably robust resource in a situation of studying intracellular infection by eukaryotic pathogens; however, a small proportion of loci undergo enough *de novo* chromatin remodeling to be detectable by ATAC-seq.

We tested whether there was a relationship between these dynamic chromatin alterations and transcriptional changes. We grouped genes according to the patterns of ATAC-seq peaks in the ±150 kb flanking their transcription start sites – those with no ATAC-seq peaks in this flanking region, those with ATAC-seq peaks only in the constitutive category, unchanged with infection, and those that gained or lost ATAC-seq peaks with infection. We performed a permutation analysis which showed that the genes changing expression were very under-represented in the group with no flanking ATAC-seq peaks, but significantly over-represented in the other two categories (Figure 3a).

**Figure 3.**
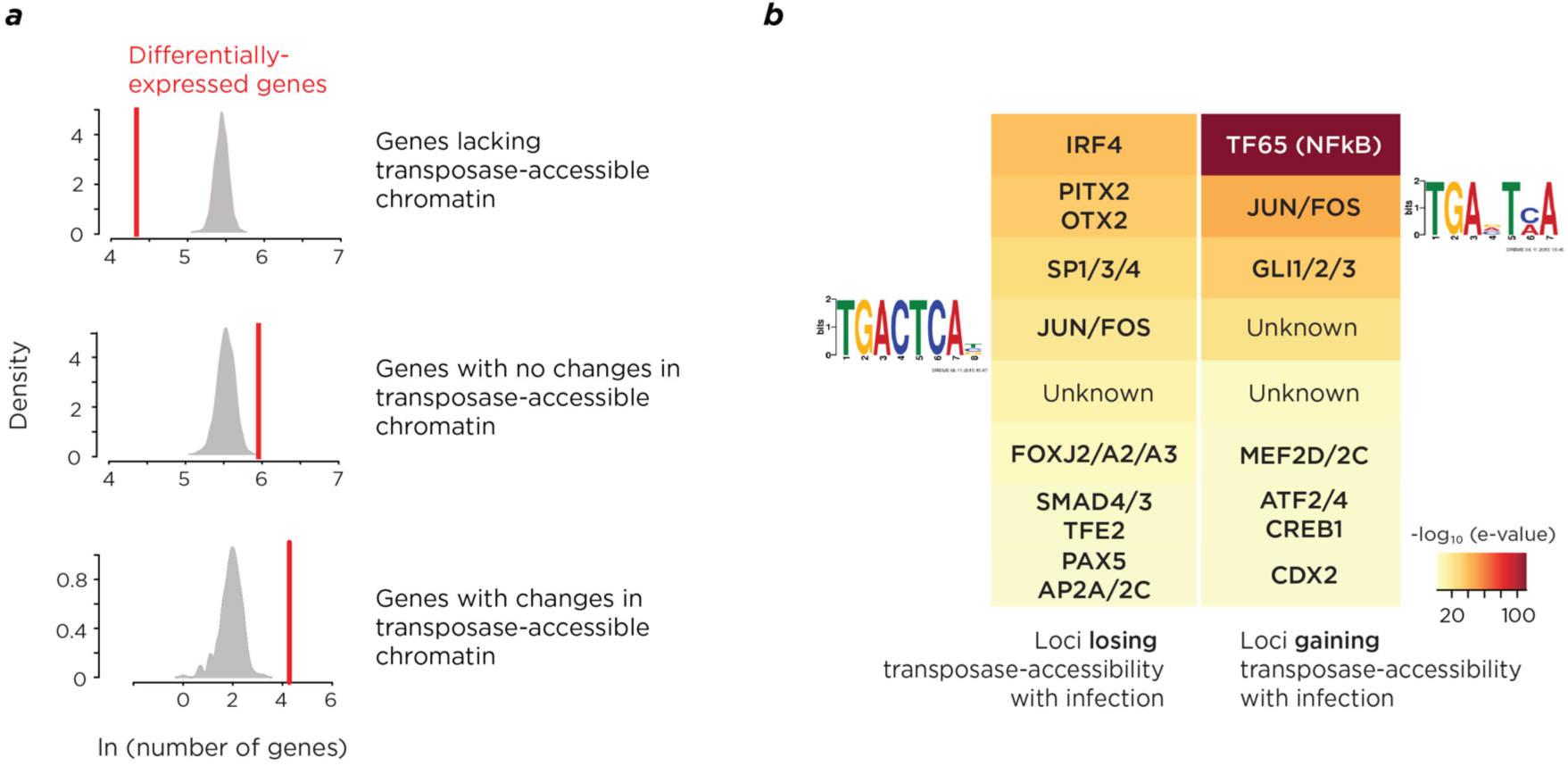
Specific patterns of host cell chromatin modifications occur with *T. gondii* infection. **(a)** Permutation studies were performed to test whether the occurrence of genes changing expression with infection was non-randomly associated with each of three categories of genes, (i) those with no loci of open, transposase-accessible chromatin within the ±150 kb flanking the transcription start site, (ii) those with flanking open chromatin where there were no changes with infection, and (iii) those with changes in flanking patterns of open chromatin. The frequency of genes with differential expression (red) is significantly enriched compared to randomly selected genes (grey) for categories (ii) and (iii), with significant depletion for category (i). In **(b)** we show a summary of motif analysis at the loci changing transposase-accessibility with infection. While some motifs could not be matched (unknown) to known transcription factors (TFs), a number of known TF binding sites were identified. Of these, the AP-1 components JUN and FOS were enriched both at loci losing and gaining transposase accessibility, supporting a model of redistribution of binding with infection.

### Specific transcription factor binding site motifs are enriched at loci changing chromatin accessibility

As transposase-accessible chromatin is likely to represent loci bound by TFs in a sequence-specific manner [37], we explored whether a motif analysis could help to identify which TFs were associated with a redistribution of *cis*-regulatory loci in infected HFFs. We tested the loci losing and gaining transposase-accessibility separately using the MEME algorithm [38] of MEME-ChIP [39]. The motifs found were then compared to known TF binding sites in the JASPER database [40]. A number of motifs were found resembling those known to be bound by mammalian transcription factors (Figure 3b). The most strongly enriched motif overall was that for TF65, the RELA/p65 subunit of NFkappaB, which has been described to be translocated into the nucleus by *T. gondii* during infection [41]. This motif is enriched specifically at loci that gain chromatin accessibility with infection, suggesting that the nuclear translocation is accompanied by *de novo* binding of the transcription factor to its target sites in the genome, with accompanying displacement of nucleosomes locally, resulting in the opening of chromatin we found using ATAC-seq.

We were especially interested in the pattern manifested by the Activator Protein-1 (AP-1, c-Jun/c-Fos) transcription factor, whose motif was enriched in both sets of loci (Figure 3b). This suggested a model of redistribution of AP-1 binding associated with *T. gondii* infection. While c-Jun has previously been described to be induced early in the response to *T. gondii* infection [10], we did not confirm any changes in the levels of transcription of *JUN* or *FOS*, as the RNA-seq data for these genes do not show significant variation after 24 hours of infection (**Figure S3**).

### *A core group of “hub genes” is targeted by* T. gondii *infection*

With two orthogonal data sets of gene expression and chromatin accessibility available to us, we asked whether we could identify the genes with the greatest effects to mediate the response to the *T. gondii* infection. We ranked the pathways derived from genes with gene expression and chromatin accessibility changes, and identified the top 20 shared pathways (**Table S2**). We made the assumption that there would be “hub genes” [42] enriched across these 20 pathways. We found 93 genes to contribute to at least 4 of these 20 pathways (**Table S3**), allowing us to focus on these as candidate hub genes, studying the interactions of their protein products using GeneMANIA [43]. We show the results of this analysis in Figure 4. The results reveal a subgroup of 10 proteins to be central to the interactions of the broader group.

**Figure 4.**
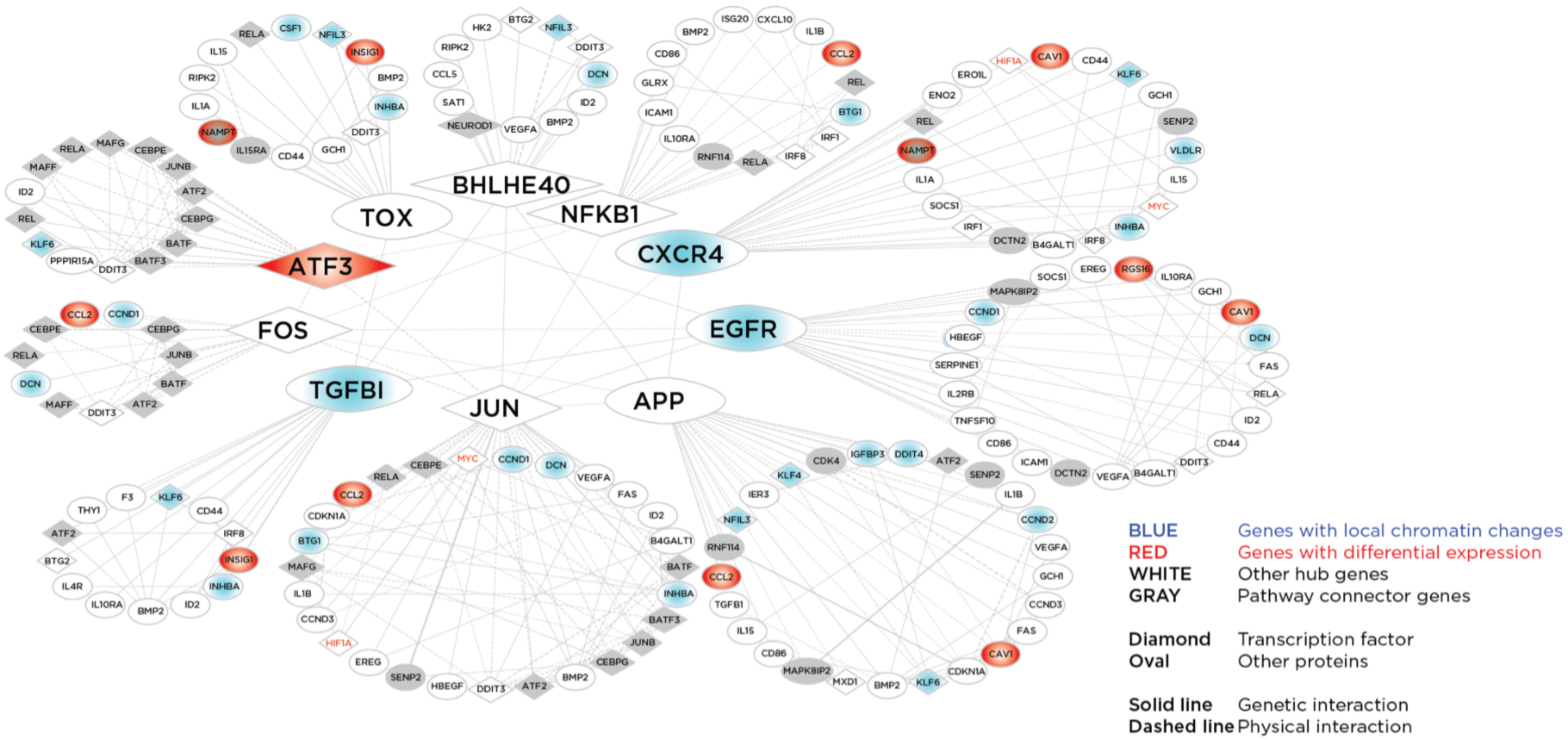
Identification of hub genes mediating the host response to *T. gondii* infection. The 93 genes contributing most frequently to the 20 pathways most commonly implicated by our transcriptomic and chromatin studies were studied for protein-protein interactions, revealing a central group of 10 genes with major effects in the host response to infection, ordered by decreasing number of interactions, starting with JUN and proceeding anti-clockwise. We find that the AP-1 components JUN and FOS are both represented as hub genes that account for the widespread transcriptional perturbations observed in *T. gondii*-infected cells.

The presence of many of these hub genes is consistent with prior observations about the host response during *T. gondii* infection. NFkB signaling is involved in the host immune response to *T. gondii* [44], while EGFR [45] and TGF-beta [46] have each previously been described to be induced during *T. gondii* infection. We note that despite the approach being methodologically independent of our motif studies, this hub gene analysis also defines the AP-1 genes *JUN* and *FOS* as central to the response to *T. gondii* infection. Neither *JUN* nor *FOS* has local changes in chromatin structure (reflected by the absence of blue shading in Figure 4), consistent with their unchanged expression levels (**Figure S3**), and indicating that their altered function is likely to be mediated by post-transcriptional mechanisms.

### Chromatin immunoprecipitation of c-Fos shows redistribution with infection

The model suggested by these ATAC-seq and gene network studies is one of redistribution of the binding sites of AP-1 in response to *T. gondii* infection. To test this, we performed chromatin immunoprecipitation (ChIP) on uninfected and infected HFF cells. We focused on loci where the ATAC-seq data indicated the acquisition of transposase-accessible chromatin only following infection (infected-only), and where the canonical 5’-TGANTCA-3’ AP-1 binding sequence was also located. We immunoprecipitated chromatin from uninfected and infected HFFs using an antibody against c-Fos, testing for enrichment using single locus quantitative PCR (qChIP). Positive and negative control loci were used to demonstrate the expected degree of enrichment for loci binding c-Fos. At the loci acquiring transposase-accessibility with infection, the qChIP results showed evidence for binding of c-Fos in the infected but not the uninfected cells (Figure 5). These results demonstrate that, within the 24 hour infection period, c-Fos (presumably in combination with c-Jun as part of the AP-1 TF) binds to new loci in the genome that are also characterized by the *de novo* acquisition of transposase accessibility.

**Figure 5.**
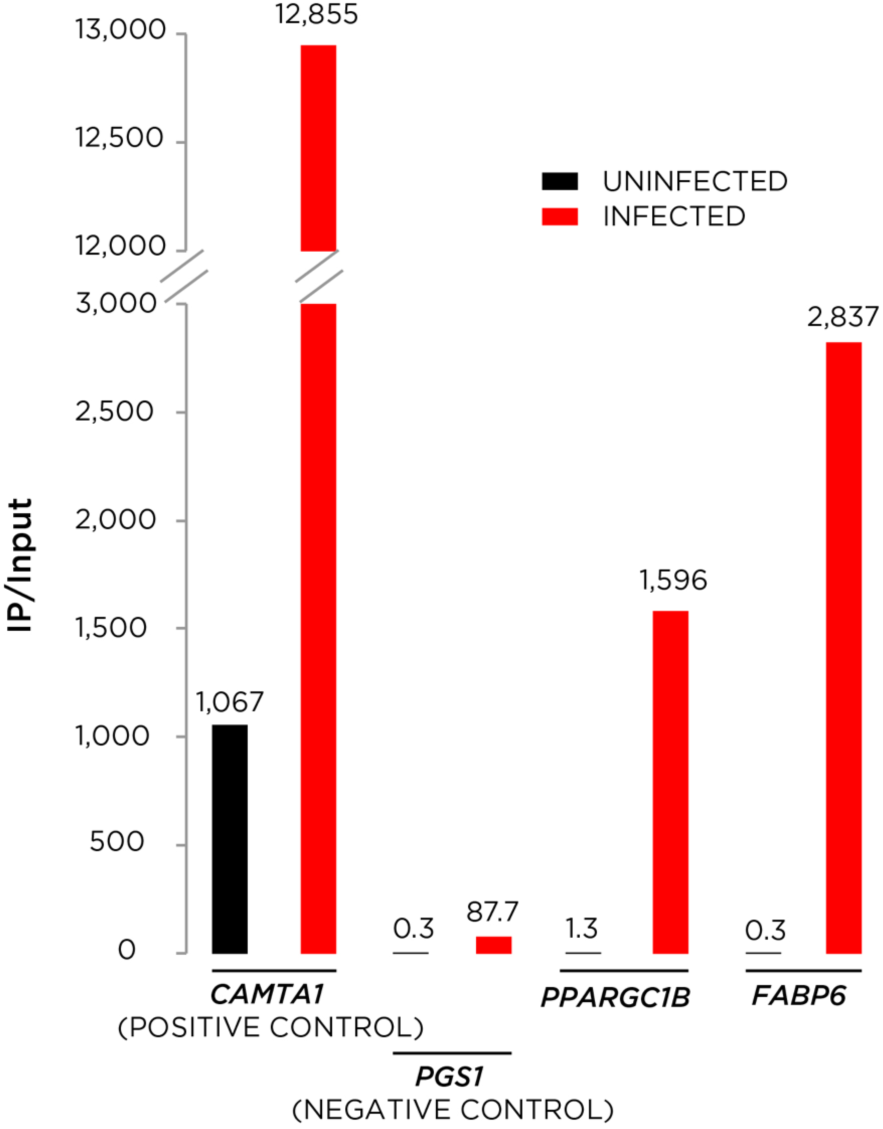
Quantitative single locus chromatin immunoprecipitation (qChIP) studies of the host cell confirms the acquisition of ectopic c-Fos binding with *T. gondii* infection. The promoter of the *CAMTA1* gene was used as a positive control, as it had constitutive transposase-accessibility in uninfected and infected cells, contains the AP-1 binding motif, and has also been found to bind c-Fos in GM12878 cells (http://www.encodeproject.org/experiments/ENCSR000EYZ/). The negative control *PGS1* locus amplicon lacks all of these characteristics, but has the intronic location comparable with the loci tested in the *PPARGC1B* and *FABP6* genes, where ATAC-seq showed loci with transposase-accessibility only in infected cells. We show the ratio of immunoprecipitated (IP) to input quantification from real time PCR with a clear enrichment in infected cells (red) at the *CAMTA1* locus (>1,000) relative to the *PGS1* locus (<100). The enrichments at *PPARGC1B* and *FABP6* also showed patterns indicating binding of c-Fos only in the infected and not the uninfected cells, lending support to the model of AP-1 redistributing to these loci with *T. gondii* infection.

## DISCUSSION

When host cells have been infected by *T. gondii*, by 24 hours, they show clear changes in gene expression that is reproducible across multiple studies, prior to undergoing cellular decompensation or death. We find that at this timepoint there is also a shift towards aerobic glycolysis, and that changes in DNA methylation and hydroxymethylation are significantly enriched at loci of transposase-accessible chromatin flanking gene promoters, but that only a very small minority of genes changing expression have such changes in their flanking regions. In contrast, the more striking finding was an infection-associated redistribution of a subset of loci of transposase-accessible chromatin, loci enriched for binding site motifs for TFs including that of AP-1 (Jun/Fos). We show that the genes with expression changes and flanking alterations of chromatin are targets of AP-1 regulation, and that chromatin immunoprecipitation for the c-Fos protein reveals alterations of its enrichment pattern concordantly with the patterns of closing and opening of chromatin associated with infection.

This study uses genome-wide approaches and an experimental design that is free from some of the major confounding variables that we have described to affect epigenetic association studies [47,48], including cell subtype proportional alterations and genetic polymorphism, as we used the same HFF cell line in all experiments. As with prior studies of intracellular pathogenic infections, we find that there are changes of DNA methylation (5(h)mC) of the host cell’s DNA [1–5], including changes of the 5hmC modification which has not previously been tested. We used the approach to focus on changes occurring within loci of accessible chromatin to generate higher-confidence findings within candidate *cis*-regulatory elements. Since a *T. gondii* infection is associated with HFFs failing to progress through the cell cycle [49], these dynamic changes in cytosine modifications are not attributable to effects associated with DNA replication, and occur within 24 hours, an unusual insight into the temporal dynamics of DNA and chromatin modifications in non-dividing cells.

We were prompted to study the metabolic state of the infected cells by the loss of 5hmC indicated by the HELP-GT assay. A mechanism for a change of 5hmC could involve the alteration of co-factor availability for the TET enzymes catalyzing the hydroxylation of 5mC to 5hmC. Alpha-ketoglutarate is a co-factor for the TET enzymes, and could be diminished in production by a switch to aerobic glycolysis in infected cells. Aerobic glycolysis is in part regulated by the activation of HIF1alpha [50], which has been found to occur in other intracellular infections including those by *Leishmania donovani* [51] and *Mycobacterium tuberculosis* [52]. We found evidence for a shift towards aerobic glycolysis in the infected HFFs, but no associated diminishment of TET enzyme activity. Mass spectrometry showed a non-significant shift towards less 5hmC in the infected cells, but we note that the DNA harvested from the infected cells contains a proportion of *T. gondii* DNA, which we have previously shown to be lacking in cytosine modifications [36]. The HELP-GT assay only tests a subset of potentially hydroxymethylated loci in the genome, so the results of the mass spectroscopy, testing cytosines throughout the genome, can be considered to more accurately reflect TET activity changes. We conclude that while the *T. gondii*-infected cells manifest a change in metabolism towards aerobic glycolysis within 24 hours of infection, this does not lead to inhibition of TET function or the loss of 5hmC.

The remodeling of chromatin in this 24 hour period is striking, with several percent of the loci of open chromatin defined by ATAC-seq lost with infection, and a comparable number of loci becoming transposase-accessible during that time. Our model for how this occurs is based on our implication of the AP-1 TF mediating the process based on both motif analysis at the loci changing chromatin accessibility, and the central role for the JUN and FOS components of AP-1 as mediators of the changes observed using hub gene analysis. The *de novo* accessibility of chromatin indicates the active displacement of nucleosomes, a property of so-called pioneer TFs [53]. AP-1 has been found to cooperate with the glucocorticoid receptor in accessing nucleosomally-occupied binding sites [54], suggesting that it could be working as one of multiple TFs involved in the chromatin remodeling during *T. gondii* infection.

The mechanism for altered TF function and activity during infection by *T. gondii* infection is not known, but may involve interaction with enzymes secreted by the parasite into the host. Dozens of GRA and ROP proteins are secreted by *T. gondii* and function as effectors in the host, with several linked to parasite virulence and changes of the host intracellular environment [11,12,55– 59]. The mechanisms of interactions of many ROP and GRA proteins with the host are increasingly recognized. GRA16 traffics to the nucleus and complexes with HAUSP and PPA phosphatase to modulate the cell cycle and the p53 pathway [11], whereas GRA24 forms a complex with host p38a and induces its activation [58] with induction of inflammatory genes, including *EGR* and *FOS*. The ROP16 kinase enters the nucleus and modulates expression of STAT signaling pathways [12], whereas expression of ROP38 kinase, which does not appear to target the host nucleus, downregulates the MAPK pathways and inflammatory gene expression, suggesting a fine tuning of parasite secreted effectors [59]. Our finding that the most strongly enriched motif in loci gaining chromatin accessibility is that for the RELA/p65 subunit of NFkB is consistent with prior reports of its translocation into the nucleus being caused by *T. gondii* [41]. Our genome-wide approaches reveal evidence for alterations in NFkB and EGFR/TGFb signaling during *T. gondii* infection, and builds upon prior observations about AP-1 induction by intracellular *T. gondii* [10,12,60], revealing AP-1 to have a central role in transcriptional changes, mediated by a systematic change in binding site locations for a subset of loci, likely to occur in partnership with other TFs, possibly influenced by new post-translational modifications induced by the parasite.

## MATERIALS AND METHODS

### Cell and parasite cultures

Cell and parasite cultures were maintained as previously described by our lab [61]. Human Foreskin Fibroblasts (HFFs) were grown in Dulbecco’s modified Eagle medium (DMEM) supplemented with 10% fetal bovine serum, 100 U/mL penicillin, 100 μg/mL streptomycin and 2 mM L-glutamine and were maintained at 37°C with 5% CO_2_. *T. gondii* Type I tachyzoites (RHΔHxΔKu80) were also maintained at 37°C with 5% CO_2_ by infecting 25 cm^2^ flasks containing confluent HFFs every 48 hours.

### Host cell infections

The 25 cm^2^ flasks containing freshly egressed *T. gondii* tachyzoites were then passaged through a 25-gauge needle three times to lyse out any remaining intracellular tachyzoites from their hosts, and the tachyzoites were spun down at 3,000 rpm for 8 minutes. 75 cm^2^ flasks containing confluent HFFs were then infected with these tachyzoites using a multiplicity of infection (MOI) of 3, and 24 hours after the infection the percentage of infected cells per flask was calculated. Flasks counted as at least 80% infected at this time were harvested by scraping. A flask containing HFFs to be used as an uninfected control was also harvested in parallel by scraping. The cells were pelleted by centrifugation at 1,300 rpm for 5 minutes. The cells were harvested such that both RNA and DNA could be extracted from the same flask for each biological replicate, and a total of three biological replicates of both the control and infected flasks were harvested in total.

### DNA extraction

Genomic DNA (gDNA) was extracted from each sample as follows: 500 μL 10% SDS and 10 μL RNase A (10 mg/mL) were added to each sample followed by an incubation at 37°C for 1 hour. 50 μL of proteinase K (20 mg/mL) was then added and the samples were incubated at 55°C overnight. An equal volume of saturated phenol was then added to each tube, and the samples were mixed slowly at room temperature for 15 minutes. The samples were then spun down at 3,000 rpm for 10 minutes and the supernatant was collected. A total of 3 phenol extractions were performed, followed by 2 chloroform extractions. The samples were then pipetted into a dialysis bag and further purified at 4°C through three changes of a 20X NaCl, sodium citrate (SSC) buffer over the course of 24 hours. Dehydration was then performed by covering each sample in a pile of polyethylene glycol (PEG) for ∼1 hour. The PEG was rinsed from each dialysis bag and the gDNA extracted from each sample was collected.

### RNA extraction

RNA was extracted using TRIzol reagent (Invitrogen Cat. # 15596018) according to the manufacturer’s instructions as follows. One milliliter of TRizol was added to each sample and incubated at room temperature for 5 minutes. A volume of 200 μL of chloroform was added to each sample and the tubes were shaken vigorously for 15 seconds and then incubated for 2-3 minutes at room temperature. The samples were then centrifuged at 12,000 rpm for 15 minutes at 4°C, and the aqueous phase of each sample were collected. Following this, 0.5 mL of 100% isopropanol was added to the aqueous phase, and the samples were incubated for 10 minutes at room temperature, followed by centrifugation at 12,000 g for 10 minutes at 4°C. The supernatant was removed from each tube and the pellet was washed in 1 mL of 75% ethanol and centrifuged at 7,500 g for 5 minutes at 4°C, and the supernatant was removed and the pellet was air dried for 5 minutes. The RNA pellet was then resuspended in RNase-free water and incubated at 55°C for 10 minutes. Following the RNA extraction, the samples were then treated with DNase to remove contaminating gDNA, and further purified using the miRNeasy Mini Kit (QIAGEN cat# 217004). The Bioanalyzer was used to assess total RNA integrity prior to library preparation, and only samples with a RNA integrity number (RIN) greater than 8 was used for further downstream library preparation.

### Directional RNA-seq library preparation

Directional RNA-seq libraries were prepared as previously described by our lab [62]. One microgram of RNA was DNase-treated and rRNA-depleted (Ribozero rRNA Removal Kit, Epicentre). RNA was used as a template for SuperScript III first-strand cDNA synthesis (ThermoFisher Scientific, SuperScript III kit cat# 18080-044), using oligo-dT as well as random hexamers. Actinomycin D was added to the reaction to prevent any possible amplification from contaminating gDNA. During second-strand synthesis, a (dUTP/dATP/dCTP/dGTP) mix was used to create directional libraries. Before library preparation, cDNA samples were Covaris-fragmented to 300 bp fragments. The samples were then end-filled, 3’ terminal A extended and ligated with TruSeq-indexed Illumina adapters. Uracil-DNA-glycosylase (UDG) treatment preceded the PCR reaction to amplify exclusively the originally oriented transcripts. Libraries were amplified using P5 and P7 Illumina primers. Prior to sequencing the libraries were gel-extracted for size selection, primer-dimers were removed and the quality was assessed on the Bioanalyzer. The six libraries were multiplexed on a single lane of the Illumina Hi-Seq 2500 platform twice to obtain 100 bp single-end reads. The reads combined from the two lanes of the sequencer for each sample were used for further downstream analysis, and this resulted in a mean of 25 million reads per sample (**Table S4**).

### RNA-seq differential expression analysis

The samples were aligned to a combined *Homo sapiens* (hg19)-*T. gondii* (Me49 version 9.0) genome. Only complete chromosomes were considered; incomplete genomic contigs from both species were excluded. We removed potential PCR duplicates from the RNA alignments using *MarkDuplicates* from *Picard Tools* (version 1.1.29). Each sample was aligned using *GSNAP* alignment software [63]. We counted reads on specific gene features (exons) using the *union* strategy within *Htseq-count* (version 0.6.1) [64]. The Bioconductor *DEseq* package [65] was used for normalization across all six samples through size factor estimates, and *DEseq* was further used for differential expression analysis by fitting a negative binomial model to our samples. The list of differentially-expressed genes is provided in **Table S5**.

### HELP-tagging/HELP-GT library preparation

HpaII enrichment by ligation-mediated PCR (HELP)-tagging libraries were prepared and analyzed as previously described [20], with HELP-GT libraries also prepared and analyzed as previously described [22]. A starting concentration of 500 ng of gDNA was used to generate each of the HELP-tagging/-GT libraries to measure the degree of methylation or hydroxymethylation at each CpG dinucleotide assayed. The gDNA was digested overnight with HpaII, MspI, or with MspI after a treatment with β-glucosyltransferase (BGT) in separate reactions. HpaII only digests unmethylated CpG dinucleotides, while MspI digests both methylated (hydroxymethylated) and unmethylated CpG dinucleotides. Following treatment with BGT, hydroxymethylated CpG dinucleotides are protected from digestion by MspI, and therefore can be identified from the methylated or unmethylated CpG dinucleotides. Following restriction enzyme digestion, the samples were purified by a phenol/chloroform extraction followed by an ethanol precipitation. After purification, the samples were ligated first to Illumina adapters containing the T7 promoter sequence and the EcoP151 recognition site (AE adapters) using a New England BioLabs Quick Ligation Kit (NEB cat #M2200S). After ligation, the DNA samples were purified using Agencourt AMpure beads and then digested with EcoP151 at 37°C overnight, end-repaired, 3’ terminal A extended and then a second Illumina adapter that contains the Illumina sequencing primer sequence was ligated (AS adapter), again using a New England BioLabs Quick Ligation Kit (NEB cat #M2200S). After ligation, the products were purified using Zymo DNA Clean and Concentrator, and then *in vitro* transcribed using the Ambion MEGAshort kit (Life Technologies). Following *in vitro* transcription, the samples were purified using an RNeasy clean-up Kit (Qiagen) before reverse transcription was performed using a Superscript III kit (ThermoFisher Scientific, SuperScript III kit cat# 18080-044). The first cDNA produced was used as a template for a PCR using the following conditions 96°C for 2 minutes, then 18 cycles of 96°C for 15 seconds and 72°C for 15 seconds followed by 5 min at 72°C for the final extension. After the PCR, the library was purified using Zymo DNA Clean and Concentrator. Prior to sequencing, library quality was assessed using a Bioanalyzer, and the six libraries were multiplexed on a single-lane of the Illumina HiSeq 2500 to generate 50bp single-end reads (**Table S4**).

### HELP-tagging/HELP-GT analyses

Following library preparation and sequencing, the DNA methylation and DNA hydroxymethylation status of each assayed CpG dinucleotide can be detected. Each sample was aligned to the *Homo sapiens* (hg19) genome through the WASP pipeline [66] using the *CASAVA* aligner from Illumina (*ELAND* version 1.7.0). The formulas arctan (HpaII counts/MspI counts) and arctan (MspI-BGT counts/MspI counts) were used at each assayed CpG locus as previously described by our lab [20], using a HFF cell type-specific MspI reference to normalize the degree of DNA methylation or DNA hydroxymethylation at each locus. Candidate differential CpGs were then obtained by fitting a linear model at each locus to estimate the significance of the difference in DNA methylation or DNA hydroxymethylation due to the infection with *T. gondii*. Loci with differential hydroxymethylation were removed prior to identifying loci with differential 5(h)mC from HELP-tagging results, to reduce the effect of 5hmC changes when identifying differential patterns of differential 5mC between conditions. Loci with differential 5(h)mC and 5hmC are listed in **Table S6**.

### Nuclear extractions and measurement of TET activity

Nuclear extractions were performed on a total of three biological replicates of uninfected and infected HFFs using the EpiQuik Nuclear Extraction Kit I (#OP-0002) following the manufacturer’s instructions. Fresh 1X PBS was added to each flask and the cells were harvested by scraping and centrifuged for 5 minutes at 1,000 rpm. Following centrifugation, the samples were resuspended in 1X NE1 buffer. Following incubation, the samples were centrifuged for 1 minute at 12,000 rpm. The cytoplasmic extract was removed from the nuclear pellet and 10 μL per 10^6^ cells of a 1:1000 dilution of the NE2 buffer was added to each sample. The extract was incubated on ice for 15 minutes with vortexing every 3 minutes. The suspension was centrifuged for 10 minutes at 14,000 rpm at 4°C, and the supernatant containing the nuclear extract for each sample was then collected. The final protein concentration of each sample was measured using the Pierce BCA protein assay kit (ThermoFisher Scientific cat #23227) as follows. A series of dilutions of known concentrations of bovine serum albumin (BSA) was generated to create a standard curve that can be assayed alongside the samples of unknown protein concentration. Twenty five microliters of each standard or unknown sample was pipetted into a 96-well plate, and then 200 μL of the working reagent was added to each well and the plate was incubated at 37°C for 30 minutes, and finally the absorbance was measured using a plate reader.

The Epigenase 5mC Hydroxylase TET Activity/Inhibition Assay Kit (Fluorometric-#P-3087) was used to measure TET activity within three biological replicates of uninfected and *T. gondii* infected HFF samples, starting with 5 μg of nuclear extract obtained from each sample. The enzymatic reaction was performed by first adding a binding solution to each well, followed by the addition of 0.5X TET Substrate to each blank and sample well, and a diluted TET assay standard to each standard curve well. The plate was sealed and incubated at 37°C for 90 minutes. Following the incubation, the binding solution was removed and each well was washed 3x with the diluted 1X wash buffer. TET assay buffer was added to each blank, standard curve and sample well at the appropriate final concentration. The plate was sealed and incubated at 37°C for 90 minutes. Following the incubation, the binding solution was removed and each well was washed three times with the diluted 1X wash buffer. The Capture Antibody was then added to each well, and the plate was sealed and incubated at RT for 60 minutes. Following the incubation, the binding solution was removed and each well was washed three times with the diluted 1X wash buffer. The Detection Antibody was then added to each well, and the plate was sealed and incubated at RT for 30 minutes. Following the incubation, the binding solution was removed and each well was washed three times with the diluted 1X wash buffer. The Enhancer Solution was then added to each well, and the plate was sealed and incubated at RT for 30 minutes. Following the incubation, the binding solution was removed and each well was washed three times with the diluted 1X wash buffer. The signal was then detected by adding the Fluorescence Development Solution to each well, followed by incubation at RT for 4 minutes. The fluorescence was then read using a microplate reader. The TET activity of each sample was calculated as ((Sample RFU – Blank RFU) / (Protein Amount Added * Incubation Time)) * 1000. Each sample was then normalized using the total cells per sample to account for the presence of *T. gondii* protein within the infected sample and TET activity was calculated using (1000 * RFU) / (Total Cells per sample * Incubation Time).

### Measurement of 5mC and 5hmC levels by mass spectrometry

Mass spectrometry to quantify cytosine, 5-methylcytosine and 5-hydroxymethylcytosine was performed as follows. One microgram of genomic DNA was first denatured by heating at 100°C. Five units of Nuclease P1 (Sigma-Aldrich, Cat # N8630, MO, USA) were added and the mixture incubated at 45°C for 1 hour. A 1/10 volume of 1 M ammonium bicarbonate and 0.002 units of venom phosphodiesterase 1 (Sigma-Aldrich, Cat # P3243, MO, USA) were added to the mixture and the incubation continued for 2 hours at 37°C. Subsequently 0.5 units of alkaline phosphatase (Invitrogen, Cat # 18009-027, CA, USA) were added and the mixture incubated for 1 hour at 37°C. Before injection into the Zorbax XDB-C18 2.1 mm × 50 mm column (1.8 μm particle size, Agilent Cat # 927700-902, CA, USA), the reactions were diluted 10-fold to dilute out the salts and the enzymes. The samples were run on an Agilent 1200 Series liquid chromatography machine in tandem with the Agilent 6410 Triple Quad Mass Spectrometer. LC separation was performed at a flow rate of 220 μl/minute. Quantification was performed using a LC-ESI-MS/MS system in the multiple reaction monitoring mode. Results are shown in **Table S7**.

### ATAC-seq library preparation

The assay for transposase accessible chromatin (ATAC)-seq libraries were prepared as previously described [33]. A total of three biological replicates each of uninfected and *T. gondii* infected HFFs were harvested at 24 hours post infection. To prepare the nuclei, we spun 75,000 cells at 500 g for 5 minutes. Each sample was washed using 50 μL of cold 1X PBS. Samples were then centrifuged at 500 g for 5 minutes. Cells were lysed in cold lysis buffer (10 mM Tris-HCL, pH 7.4, 10 mM NaCl, 3 mM MgCl_2_ and 0.1% IGEPAL CA-630). Immediately following lysis, nuclei were spun at 500 g for 10 minutes at 4°C. The pellet was then resuspended in the transposase reaction mix (25 μL 2x TD buffer, 2.5 μL transposase and 22.5 μL nuclease-free water).

The transposition reaction was carried out for 30 minutes at 37°C and the samples were then purified using a Zymo DNA Clean and Concentrator purification kit. Following the purification, we amplified the libraries with the following PCR conditions: 72°C for 5 minutes; 98°C for 30 seconds; and a total of 10 cycles of 98°C for 10 seconds; 63°C for 30 seconds; 1 minute for 72°C. Libraries were purified using Agencourt AMpure beads. Libraries were further amplified using the PCR conditions: 98°C for 30 seconds; and a total of 14 cycles of 98°C for 10 seconds; 63°C for 30 seconds and 72°C for 1 minute. The six ATAC-seq samples were multiplexed on a single lane of the Illumina Hi-Seq 2500 to obtain 100 bp paired-end reads, which resulted in a mean of 30 million reads per sample (**Table S4**).

### ATAC-seq peak calling and motif analysis

Sequenced reads were aligned to a combined *Homo sapiens* (hg19)-*T. gondii* (ME49 version 9.0) genome using Burrows-Wheeler Aligner (BWA-MEM version 0.10) [67]. Peaks were called using MACS2 (version 2.1.0) [68] and we detected a mean of 150,000 peaks per sample (**Table S8**).

DNA motifs were discovered within the regions of transposase-accessible chromatin using the MEME algorithm [38] of MEME-ChIP [39]. The discovered motifs were compared to known TF binding sites through TomTom to match the discovered motifs to known models in the JASPER database [40].

### Integrative analyses

ATAC-seq peaks were assigned to their closest gene using the *bedtools* suite (version 2.24.0) [69], and using our RNA-seq dataset to identify the genes expressed in HFFs that had ATAC-seq peak falling within ±150 kb of that gene’s TSS. Genes were defined as having “differential” peaks if they either lost or gained ATAC-seq peaks during the infection in all three biological replicates tested, genes were defined as having “constitutive” peaks if they only had ATAC-seq peaks that were present in both the infected and uninfected samples, and genes were defined as having “no peaks” if there were no ATAC-seq peaks falling within the ±150 kb region flanking the gene’s TSS.

The total number of observed genes in each category was calculated using our list of differentially expressed genes with ATAC-seq peaks, and plotted as a ratio over the total number of expected genes calculated using all expressed genes with ATAC-seq peaks. A permutation analyses was performed using 1,000 iterations using the same number of random genes from our RNA-seq dataset to test whether the enrichment we detected was happening by random chance. The enrichment of TF binding sites within the differential regions was then identified as described using MEME-ChIP software [39]. The bedtools suite (version 2.24.0) [69] was then again used to identify the frequency of occurrence of HELP-tagging/-GT CpG loci falling within the constitutive or differential ATAC-seq peaks. A permutation analyses was again performed, again with 1,000 iterations, using the same number of a random set of CpG loci from our HELP-tagging/-GT datasets to test whether the differential CpG loci were falling within the constitutive peaks by random chance.

Potential cellular functions of the differentially expressed host genes were explored through a gene set over-representation analysis (ORA) [19] using MSigDBs Hallmark gene sets [70]. An ORA tests for enrichment within each gene set using a hypergeometric test. A p-value representing the significance of enrichment for a specific test group within each Hallmark gene set was obtained. Overlaps were computed using HUGO gene symbols and the p-value was corrected for multiple hypotheses testing using the Benjamini-Hochberg procedure to adjust the false discovery rate (FDR). We considered pathways having an FDR < 0.05 as being significantly enriched for within our list of differentially expressed genes.

A hypergeometric test was used to obtain a p-value representing the significance of the enrichment of a particular test group within each Hallmark gene set, where here “k” represents the number of genes from the test group that overlap the genes in a particular Hallmark gene set, “K” represents the total number of genes in that Hallmark gene set, “N” represents all known genes, and “n” represents the total number of genes in the test group. Overlaps were computed using HUGO gene symbols and the p-value was corrected for multiple hypotheses testing using the Benjamini-Hochberg adjustment of the false discovery rate (FDR). The significantly enriched pathways were then ranked based on the significance of their FDR, and the overlapping pathways between the test groups containing the genes with differential ATAC-seq peaks and the differentially expressed genes were identified. The rank based on the FDR of each of the overlapping pathways was then summed to obtain a new rank for each significantly enriched pathway, and then the top 20 significantly enriched pathways overlapping between the genes with differential ATAC-seq peaks and the differentially expressed genes were used to identify the hub transcriptional regulators present during the infection, and to generate the interactome network.

### Interactome network analysis

The network design was based on genetic (gene-gene) and physical (protein-protein) relations made using *Cytoscape* (version 3.2.1) [71]. For this purpose we used the *GeneMANIA* (version 3.4.0) plugin for *Homo sapiens* interactome data to analyze the list of hub transcriptional regulators [72], and the ontological analyses to identify any TFs were performed using the *BiNGO* module. The interaction network was then integrated with gene-signaling pathway information obtained from the GSEA database [70]. For the initial visualization, the results were sorted by the circular layout option, which sorted nodes on the basis of the number of interactions between them. These results allowed the identification of the genes with the highest number of interactions with other genes and were associated with the significantly enriched signaling pathways.

### Chromatin immunoprecipitation

The Myers lab chIP-seq protocol (v. 041610.1) was optimized for use on HFFs as follows. A total of 4 × 10^7^ HFFs were grown to confluency in 150 mm^2^ dishes. These dishes were infected with *T. gondii* as described, cross-linked and harvested 24 hours post infection. 4 × 10^7^ HFFs were also cross-linked and harvested in parallel as an uninfected control. The infected and uninfected dishes were cross-linked as follows: formaldehyde was added directly to the media to a final concentration of 1% and the dishes were then incubated for 10 minutes at room temperature. To stop the cross-linking reaction, glycine was added to a final concentration of 0.125 M. The cells were then washed with cold 1X PBS and scraped off the dishes and transferred into 15 mL conical tubes on ice and pelleted by centrifugation at 2,000 rpm for 5 minutes at 4°C.

Each cell pellet containing 20 × 10^6^ cells was resuspended in 1 mL Farharm Lysis Buffer. Samples were then pelleted by centrifuging at 2,000 rpm for 5 minutes at 4°C and resuspended in 300 μL cold RIPA buffer. The samples were then processed in a Biorupter using the high setting for a total of 20 minutes with 30 seconds on and 30 seconds off. An average shearing size of 200 bp was obtained for each sample. The sonicated mixture was then spun at 14,000 rpm for 15 minutes at 4°C and the supernatant was collected. Fifty microliters of sheep anti-rabbit magnetic Dynabeads (ThermoFisher cat#11203D) was added to a 2 mL tube containing 1 mL PBS/BSA. Tubes were washed 3x and 6 μg of anti-cFOS antibody (UC Santa Cruz cat# sc-7202) was added to each tube. The antibody was coupled to the beads overnight at 4°C. Antibody-coupled beads were then washed again 3x with PBS/BSA and then added to the 1 mL chromatin preparation obtained from 4 × 10^7^ cells and again incubated overnight at 4°C. The beads containing the immuno-bound chromatin were collected using a magnet, followed by first washing 5x in cold LiCl wash buffer and then washing 1x in cold 1X TE buffer. The supernatant was then discarded and the bead pellet was resuspended in 200 μL IP elution buffer at room temperature.

The samples were incubated at 65°C for 1 hour and vortexed every 15 minutes to elute the immuno-bound chromatin from the beads. The samples were then spun at 14,000 rpm for 3 minutes at room temperature. The supernatant containing the immunoprecipitated DNA was incubated overnight at 65°C to complete the reversal of the formaldehyde cross-links and then purified using the Zymo purification kit.

The publically available c-FOS ChIP-seq data for the GM12878 cell line obtained through the ENCODE project was used to identify a positive locus that could be used to test our immunoprecipitation prior to sequencing. The c-FOS peaks obtained for the GM12878 cell line, along with our own ATAC-seq HFF peaks were uploaded to the UCSC genome browser (https://genome.ucsc.edu/). We identified regions that contained ATAC-seq peaks within all six of our HFF samples (constitutive peaks present between the uninfected and infected HFFs), as well as a c-FOS GM12878 peak and the “TGANTCA” c-FOS binding motif identified through the MEME-ChIP analysis. We designed our PCR primers flanking the “TGANTCA” motif. A negative locus was chosen based on the absence of the above-described marks. Prior to library preparation and sequencing, 0.5 μL of each IP was used for qPCR in duplicate to test for the enrichment of the positive locus, after normalization to an input control. All PCR primers used in this project are shown in **Table S9**.

## ACKNOWLEDGEMENTS

We thank Drs. Kami Kim and Inessa Gendlina for guiding the *T. gondii* experimental studies performed. NU was supported by NIH training grant T32 NS007098. NAW and ADJ were supported by the Medical Student Training Program (NIH T32 GM007288). Einstein core facilities involved were the High-Performance Computing Core, the Epigenomics Shared Facility, and the Genomics Core Facility, with support from the Albert Einstein Cancer Center (P30CA013330) and the Center for Epigenomics.

## PUBLIC REPOSITORIES

Software: GitHub

https://github.com/GreallyLab/Ulahannan-et-al.-2016

Sequence data: GEO Accession number: GSE79612

http://www.ncbi.nlm.nih.gov/geo/query/acc.cgi?acc=GSE79612

